# Establishing CREATE: Lessons learned in setting up a training environment for early-career researchers in respiratory medicine

**DOI:** 10.1101/2021.10.20.465179

**Authors:** Katherine Christian, Alison Hey-Cunningham, Tamera Corte, Nicole Goh, Jade Jaffar, Paul Reynolds, Alan Teoh, Lauren Troy

**Affiliations:** NHMRC Centre of Research Excellence in Pulmonary Fibrosis, Australia; The University of Sydney Central Clinical School, Sydney, Australia; Royal Prince Alfred Hospital, Camperdown, Australia; Austin Health and Alfred Health, Melbourne, Australia; Faculty of Medicine, University of Melbourne, Australia; Department of Allergy, Immunology and Respiratory Medicine, The Alfred Hospital, Melbourne, Australia; Monash University, Melbourne, Australia; Lung Research Laboratory, Faculty of Health and Medical Sciences, University of Adelaide, Australia; Department of Thoracic Medicine, Royal Adelaide Hospital, Australia

**Keywords:** Early-career, professional development, Australia, training, health and medical research

## Abstract

**Background:** The purpose of the National Health and Medical Research Council Centre of Research Excellence in Pulmonary Fibrosis (CRE-PF) is to improve and extend the lives of patients living with pulmonary fibrosis through the development of a comprehensive and integrated program of basic and clinical research and education across Australia. A key objective of the CRE-PF was establishment of a unique national training scheme, CREATE, for early-career researchers (ECRs) in respiratory research. CREATE ECRs are broadly drawn from two main fields of researchers: clinicians and scientists, where clinicians tend to be involved in part-time translational research and scientists are involved in broad scientific research including laboratory or genetic research, health economics or population research.

**Methods:** We describe the CREATE Program which, with limited budget and the assistance of key organisations, has provided funding opportunities (scholarships, fellowships, prizes, travel and collaboration grants), professional development (mentoring program, symposia, presentation opportunities and on-line training) and fostered a connected, supportive research community for respiratory ECRs.

**Results:** The CREATE program has successfully fostered the development of the supported researchers, contributing substantially to the future of pulmonary fibrosis research in Australia. During the life of the program the CRE-PF has offered 10 PhD scholarships and five postdoctoral fellowships, awarded 13 travel grants and three grants to promote collaboration between ECRs from different institutes. A mentoring program has been established and CREATE Symposia have been held in association with key meetings. During COVID-19 restrictions, a series of virtual research meetings has offered 12 CREATE ECRs from seven universities the opportunity to present their research to a national audience.

CREATE research-related achievements are impressive, including over 80 first-author publications by ECRs, and many conference presentations. Contributions to the research community, measured by committee membership, is also strong.

**Conclusions:** In spite of a very limited budget, wide geographic distribution of participants and the multi-disciplinary nature of the cohort, we have succeeded in providing a unique, supportive academic development environment for CREATE ECRs. Lessons learned in the process of developing this program include the importance of leveraging funding, being flexible, building networks and seeking and responding to ECR input.

## Background

The purpose of the National Health and Medical Research Council Centre of Research Excellence in Pulmonary Fibrosis (CRE-PF) is to improve and extend the lives of patients living with pulmonary fibrosis through the development of a comprehensive and integrated program of basic and clinical research and education across Australia.

A key objective was establishment of a national training scheme for respiratory researchers. The CRE-PF vision was to train early-career researchers (ECRs), known in our program as CREATE (<u>CRE A</u>dvanced <u>T</u>raining <u>E</u>nvironment) Fellows, to contribute to the research workforce. This would be achieved by building and translating knowledge from bench-to-bedside with a focus on the prevention of end-stage fibrotic lung disease and the development of new treatments to improve the health of Australians affected by pulmonary fibrosis. CREATE ECRs are broadly drawn from two main fields of researchers: clinicians and scientists, where clinicians tend to be involved in part-time translational research and scientists are involved in broad scientific research including laboratory or genetic research, health economics or population research.

Clinician researchers represent a unique niche in health and medicine, bridging the gap between scientific discovery and patient care (1,2); and, although very highly valued, clinical researchers are unfortunately increasingly rare (3). At the same time, science postdoctoral fellows are often inadequately linked to clinical colleagues, meaning that the “relevance” of their work may be seen as obscure to practising physicians and indeed the public (4). Health and medical research needs composite teams of clinical and non-clinical researchers to enable continued progress (2).

In recent times it has been recognised that there needs to be more emphasis on holistic researcher development (5). Conventional training for researchers in (STEMM) disciplines in Australia and internationally, has been found wanting (6,7). Publishing and disseminating research results, building networks at scientific conferences, and securing grant funding are the components that have been found to contribute to a successful track record and ability to attract grant funding (8).

The aim of the innovative CREATE training scheme has been to expand “conventional” training for emerging leaders by including key aspects of career development through both experiential and theoretical learning and by providing opportunities to broaden their networks within the field and build their research track record. A successful postdoctoral culture needs to provide ECRs with both productivity skills and a broader “village” that becomes their supportive framework (9). It also needs to teach ECRs not only research methodology, how to *do* research but also how to build the track record necessary for a research career (10). Focus of the evidence-based CREATE program has included provision of recommended initiatives including funding opportunities for ECRs (11), multi-faceted professional development (12) and fostering of the supportive framework ECRs need to thrive (9). The program encompasses establishing collaborations to ensure translational relevance; connecting to the commercial research environment; strategic publishing; communication with diverse audiences including policy makers and the public; budget management; expanding networks; seeking research funding and a structured mentoring element. In expanding the training available for these ECRs and providing mentoring and collaborative opportunities, our program aims to develop a generation of ECRs who will ultimately teach and mentor succeeding generations of investigators.

There have been many academic development lessons learned in the process of establishing the CREATE program. Through establishing and refining this program, our experience supports recent recognition of the importance of continual engagement between ECRs and other stakeholders, responding flexibly to the needs of ECRs, leveraging funding and providing support to build networks and professional skills.

## Methods

The CREATE Program has been developed across multiple platforms, including provision of funding opportunities, professional development activities and network building. Key to its success has been the wide-ranging collaborative support of the peak bodies in respiratory health, Lung Foundation Australia (LFA) and Thoracic Society of Australia and New Zealand (TSANZ). All aspects are overseen by the Program Advisory Committee (PAC) which ensures the CREATE program objectives are met, reviews the budget to determine activities we are able to support, and considers ways to leverage additional funding. The PAC has evolved to comprise key opinion leaders (two CRE-PF steering committee members, two ECR representatives (one clinical and one scientist), two mentor representatives with special interest in academic development (at different career stages), and management/administrative support. The PAC meets at least quarterly with ad hoc meetings as required.

CREATE was launched in 2017 with 10 ECRs, and numbers have grown rapidly. To date CREATE has 30 ECRs, 19 PhD students and 11 postdoctoral researchers. Of these 16 are medical practitioners involved in research, five others are clinician researchers in physiotherapy or psychology; the remaining eight are scientists. Each ECR has a multi-member supervisory team and, separately, most have a CREATE mentor.

CREATE ECRs must be engaged in pulmonary fibrosis research and be no more than 10 years postdoctoral (adjusted for career disruption). Prospective CREATE ECRs may be put forward by Investigators of the CRE-PF, or other leaders in pulmonary fibrosis research; applications are reviewed by the PAC. Almost all who started in the CREATE program have stayed active within it. In four years, six PhD students have finished their studies; all except one who has left research and another who has become a full-time clinician, and all postdocs, have continued within the program as CREATE Fellows. One ECR has moved beyond eligibility (as determined by years postdoctoral), electing to stay within the program as an alumna available to assist new ECRs.

Activities undertaken to fulfil the aims of the CREATE Program, outcomes of these aims and lessons learned to date are detailed below, within Results.

## Results

### Provision of Funding Opportunities

Funding for ECRs in medical research in Australia is highly competitive (13). The CREATE program has sought to offer a range of funding opportunities to encourage participation of ECRs in pulmonary fibrosis research, boost research output and help develop competitive research track records. This has included funding scholarships and fellowships, travel grants and prizes for achievement detailed below. It has become clear that even minor funding investments make a very welcome difference to these ECRs; we have partnered with LFA and industry to leverage and expand the pool of available funding.

#### Stipends and Fellowships

Attracting and retaining ECRs to the field of pulmonary fibrosis research, is integral to the CRE-PF. To this end, funding for PhD scholarships and fellowships for postdoctoral researchers were included in the successful CRE-PF grant application. The budget awarded included funding for four, three-year PhD scholarships and partial funding for three postdoctoral fellowships over two or three years for positions under the supervision of CRE-PF investigators. The initial funding was successfully extended with matching funding from the students’ institutions or by encouraging recipients to apply for their own funding once they had some initial results, thus strengthening their CVs and building their track records.

Additional monies have been secured through philanthropic donations and industry contributions received via the CRE-PF’s collaborative partner LFA. Ultimately the initial funding became ten PhD scholarships and five postdoctoral fellowships over the first four years of the program, with applications for new scholarships independently reviewed with the assistance of TSANZ. Other CREATE ECRs joined the program with funding already secured from external sources. Working closely with partners and active fundraising have helped the program to extend stipend and fellowship support to a growing number of ECRs. In 2022 the CRE-PF expects to offer two new Fellowships for CREATE ECRs to continue their research careers after completion of PhDs.

#### Travel Scholarships

Access to funding for travel and conferences is typically difficult for ECRs (11), despite conference attendance being of known importance to career development (8). Each CREATE ECR has been offered the opportunity of competitive travel scholarships to cover the costs of travel for professional development. This funding could be used for meeting attendance (including registration), or for travel to spend time with collaborators or mentors to develop skills and networks. Funding is prioritised for PhD students to encourage attendance at research meetings or for short-term international placements.

To date, 13 travel grants have been awarded; 11 for Australian conference attendance and two for travel overseas. When travel was prohibited by the COVID-19 pandemic, funding which had been allocated for travel was re-purposed for collaboration grants which are described below.

#### Collaboration Grants

In addition to the challenges posed by the general shortage of research funding, ECRs often feel they face barriers to collaboration or conducting interdisciplinary research. Barriers to collaboration include poor researcher connectivity; time constraints and physical distance; different terminology between disciplines; hierarchy; and competition (14). In 2020 and 2021, grants-in-aid were offered to specifically support new collaborations for pulmonary fibrosis research amongst CREATE ECRs from two or more institutions. These collaborative grants were offered to develop experience in building collaborations, share Australian pulmonary fibrosis resources and promote pulmonary fibrosis research opportunities within Australia. These small grants offered funding for travel or consumables or other initial project costs to break down these barriers and aid in the establishment of investigator-led research for ECRs only. Research funding in Australia is extremely competitive; short-term contracts and the ever-present need to apply for grants leads to uncertain futures for ECRs in research and high job insecurity (6,15). In most research funding schemes, ECRs must compete with more senior researchers in funding schemes with a low success rate, so it is vital for them to build a strong track record as soon as possible (8,11,16). CREATE grants were awarded expecting that this initial research would provide essential track record and preliminary findings to increase the chance of success for applications to national funding schemes.

Multiple applications were received for each round; one grant was awarded for the 2020 round, then two grants in 2021. Table 1 shows the 2020 grant-in-aid established a collaboration for two researchers from different Melbourne institutions; the two successful 2021 grants respectively brought together four applicants from three states and three applicants from three different institutions in one city.

**Table 1:**
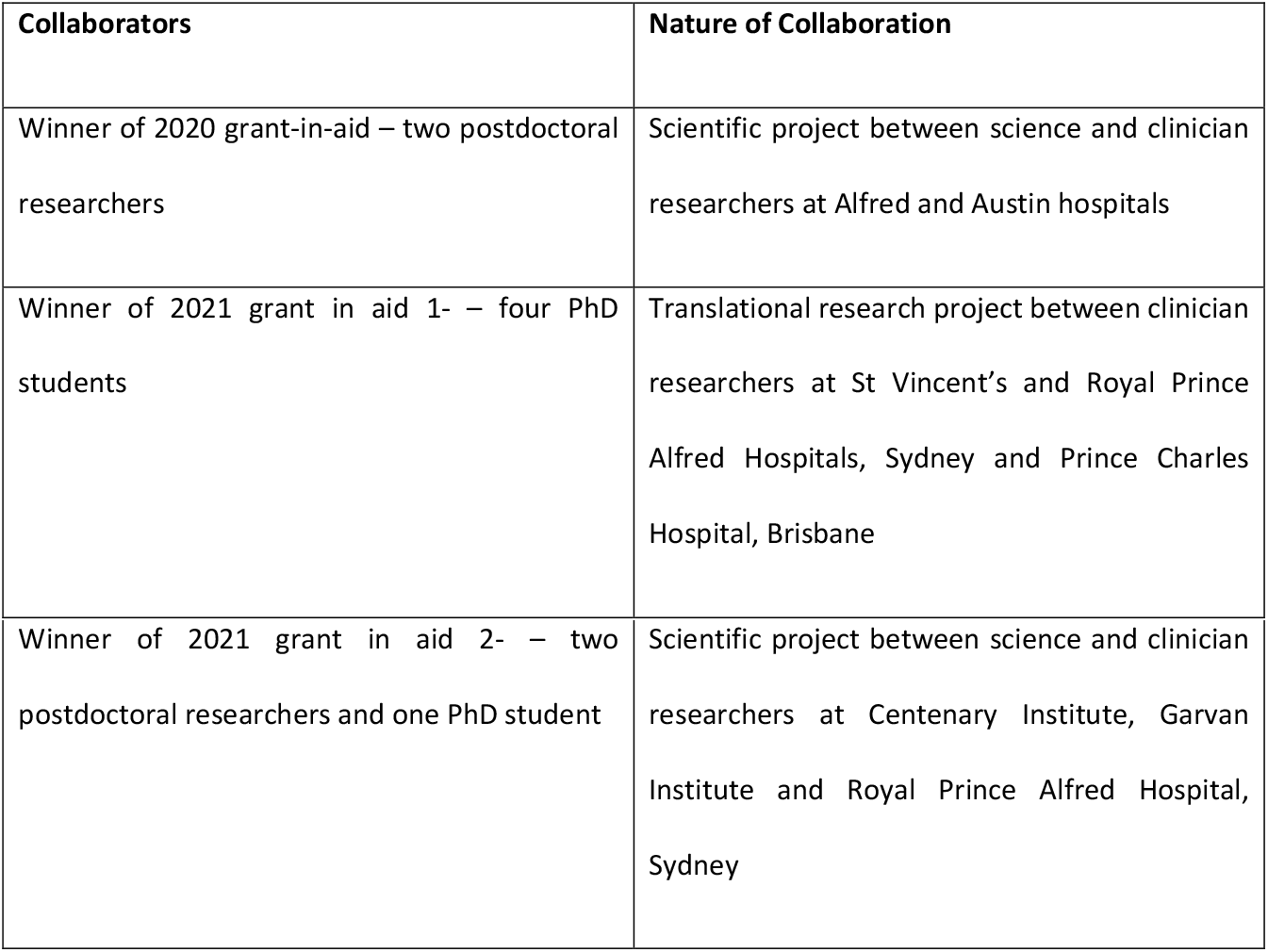
CREATE ECR collaborations funded through CREATE Grants-in-Aid.

| Collaborators | Nature of Collaboration |
| --- | --- |
| Winner of 2020 grant-in-aid – two postdoctoral researchers | Scientific project between science and clinician researchers at Alfred and Austin hospitals |
| Winner of 2021 grant in aid 1- – four PhD students | Translational research project between clinician researchers at St Vincent's and Royal Prince Alfred Hospitals, Sydney and Prince Charles Hospital, Brisbane |
| Winner of 2021 grant in aid 2- – two postdoctoral researchers and one PhD student | Scientific project between science and clinician researchers at Centenary Institute, Garvan Institute and Royal Prince Alfred Hospital, Sydney |

#### Poster Prizes

Conference poster presentations are a common way of communicating research, building profiles and adding to research track records for ECRs in many scientific and health-related disciplines. Preparation and presentation of a poster is complex, and involves far more than formatting issues and juggling images and text; it provides a way of learning to communicate research well and is part of academic apprenticeship in many health disciplines (17). Special poster prizes have been offered for CREATE ECRs as a means of encouraging them to do more than simply attend key meetings. Instead, they have actively disseminated and promoted their research, and research excellence has been recognised. Although small, these prizes for CREATE ECRs have been awarded within and alongside the usual poster award schemes at key research meetings. Three such prizes have been offered with two to come in 2021. In each case the CREATE prizes have successfully drawn attention to both the CREATE program and to the work of the individual researchers.

### Professional Development and Fostering a Supportive Network

Bringing people together, whether only CREATE ECRs or ECRs along with their more senior colleagues, has been a key strategic goal for this group. CREATE has worked within a limited budget to increasingly prioritise this as the program has developed. Professional development and networking opportunities as recommended by specialists in academic development (9,12,18) and those with a particular interest in ECRs in health and medical research (19,20) have been offered in via a mentoring program, conferences and symposia, presentation opportunities, and on-line training, as described below.

#### Mentoring Program

Although unquestionably of great value (21,22), mentoring (as opposed to supervision by one’s immediate superiors) is not available to all ECRs and formal mentoring programs within institutions are uncommon (6,23). The Franklin Women Mentoring Program, which began in 2017, was the first cross-institutional program structured mentoring program to be delivered in the health and medical research sector in Australia (24). An important and innovative aspect of the CREATE program is its mentoring system that facilitates career development. Each CREATE ECR is offered the opportunity to engage with a mentor from a different institution. The mentors are drawn from the network of CRE-PF researchers and provide the ECRs with supportive advice, honest and unbiased feedback on issues relating to their professional development. Thus, this mentorship does not conflict with the important role of the research supervisor, while expanding existing support offered to ECRs at their own Institution and laboratory group.

Initially the mentoring program was relatively unstructured outside of mentee-mentor matching. Feedback from the group indicated that a guided structure was needed to facilitate productive mentoring relationships. In response to this feedback, our approach developed such that mentees are encouraged to meet monthly with their mentors over a twelve-month period to assess development needs, discuss challenges and opportunities. Guides for both mentors and mentees have been developed as well as a suggested meeting timetable with circulation of reminders, agenda and a list of suitable topics for discussion which address commonly identified areas of need.

In addition to these regular phone or video calls, it has been encouraged that the mentor-mentee pair would typically meet in person once or twice per year at CREATE networking events held in association with major discipline meetings. Unfortunately, this was not possible in 2020-2021 during the COVID-19 pandemic.

In 2021, there were 18 active mentor pairs. An additional five mentoring relationships had formally finished (although they may, of course, continue by mutual agreement). Feedback on the program, regularly sought by email from both mentees and mentors, has overall been very positive. For example, in early 2021, 88% of mentee and mentor email respondents indicated they were satisfied or very satisfied with their CREATE mentoring experience.

Mentees who engaged with their mentors have found the experience very beneficial, as exemplified by the comments below:

> *I will definitely keep in touch MENTOR. You have definitely given me some pearls of wisdom that still ring in the back of my head when I am writing*
>
> *I am very happy with this mentoring program*

Early lessons learned from the mentoring program showed that it was best not to ask for suggestions from the mentees for their mentors, as mentors with the highest profile were over-selected. Instead it was found it was best to allocate mentors, carefully matching mentees with mentors, of any gender, from different organisations and research areas, following the example of Franklin Women (24). Initially, mentees were often shy about making initial contact, as has been experienced elsewhere (23). Unfortunately, early in the program some did not take advantage of the opportunity, whether because they were “too busy”, fearful of intruding on the mentor’s time or because they simply did not know how to go about the process. Consequently, the process has been modified to include tighter and more prescriptive recommendations for meeting formats and timing, and initiation of regular meeting reminders to mentee-mentor pairs.

#### Networking gatherings/Symposia

To achieve the CREATE objectives, certain activities have been held in conjunction with regular key meetings in the field. CREATE symposia have included dinners, a breakfast meeting and poster presentations judged by respected international researchers.

Consistent with recommendations regarding the benefit of a range of networking activities with peers and with senior researchers (19,20) these CREATE Symposia have provided the ECRs with the opportunity to meet, share ideas and showcase their work and have offered the opportunity for all involved in the CRE-PF network to meet with informal time for discussion. CREATE ECRs have also had the opportunity to meet international leaders in pulmonary fibrosis research attending the larger meeting. International leaders have taken active part in the CREATE proceedings, whether giving talks or judging posters. The CREATE budget has also contributed to travel costs for leading international researchers to come to the meetings and spend valuable face-to face time with the ECRs.

#### Virtual Research Meetings

Face to face symposia were not possible in 2020 and early 2021 due to the COVID-19 pandemic, with postponement or cancellation of the in-person components of all relevant respiratory medicine and science conferences. To promote ECR research and facilitate collaboration in the absence of in-person meetings, in mid-2020 CREATE established monthly hour-long virtual research presentations, open to all involved in pulmonary fibrosis research or treatment across Australasia. To date, virtual research meetings have been hosted by research teams from seven universities across Australia. Each key presentation has been made by a CREATE ECR, introduced by their team leader who has described their team’s overall research program. Attendance has been excellent (usually 25-30 participants); sessions have been recorded and are maintained as educational resources for the research community.

#### WhatsApp group

A CREATE WhatsApp group was established during 2019, for all CREATE ECRs, CRE-PF investigators and CREATE mentors. This replaced a Facebook group once it became apparent that this was not a preferred way for the group to connect. The WhatsApp group is a platform to share pulmonary fibrosis research news, promote relevant opportunities and successes, raise questions and seek advice, as well as to foster a sense of belonging to a network. Engagement in this group has been strong, with over 400 posts from more than 20 group members describing newsworthy research, sharing exciting presentations at research meetings and celebrating winners of awards, offering support and encouragement to the community during difficult times and sharing simple good wishes to the team. The WhatsApp group has facilitated a high level of worthwhile information exchange and fostered group connection among ECRs and between ECRs and more senior researchers.

**Figure1.**
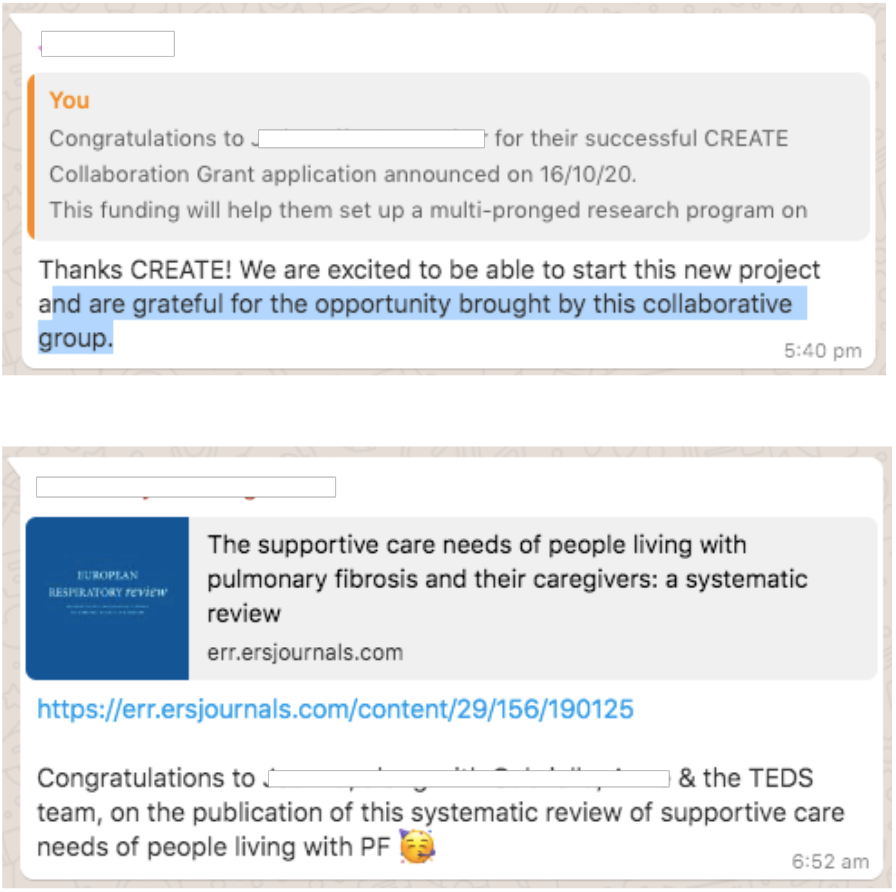
Two Examples of WhatsApp Group Messages.

#### CREATE Researcher Development in Pulmonary Fibrosis Weekend

A residential Researcher Development weekend was planned for 2020, to bring together all CREATE ECRs and their mentors from across Australia. Sponsorship was secured from a number of industry sources. Unfortunately, despite the enthusiastic response from prospective attendees, this weekend was postponed due to COVID-19. The CRE-PF has had to be flexible with the ever-changing situation. Feedback from the group indicated a strong desire to wait and hold an in-person event, now planned for November 2021. The weekend will offer intensive researcher development on topics recommended by organisations such as Vitae in the UK (25,26) and the Council of Graduate Schools in the USA (19). This will include educational and upskilling sessions on topics selected by the ECRs as their areas of need, along with networking and collaboration building opportunities; and team building experiences. With generous support from industry, all costs for the ECRs will be covered, removing financial barriers to participation. See program overview in Supplementary table.

#### Online learning

Early in the CREATE program, a series of online modules was developed and made available through a secure section of the CREATE website. These modules were designed to offer generic management training on a range of topics identified as specific ECR needs and can be completed at any time. The CREATE online modules shown in Table 2 were designed to enhance and complement the training and guidance our ECRs received locally.

**Table 2:**
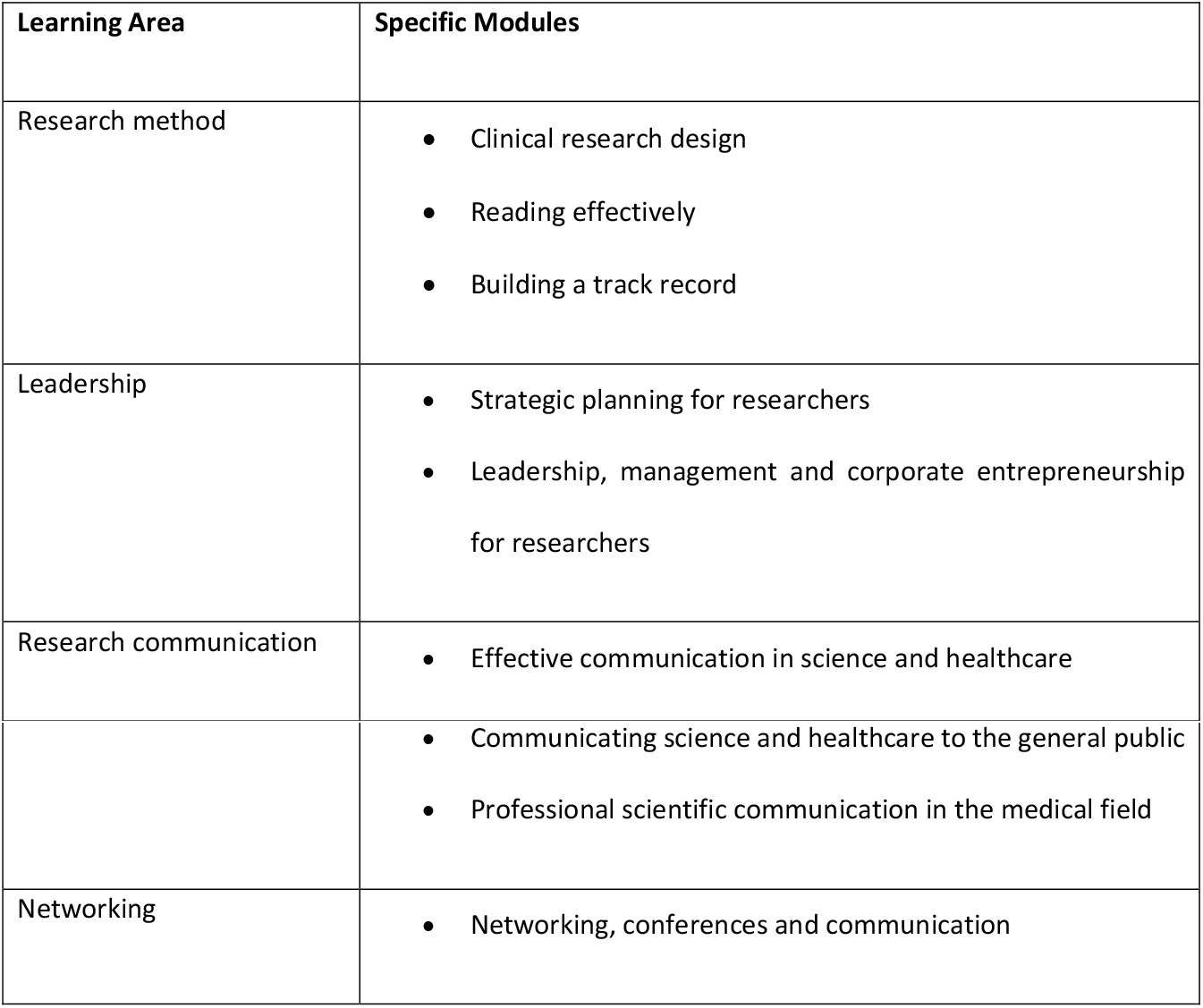
Online Learning Modules for CREATE ECRs.

Uptake of the modules has been disappointing. Unfortunately, this is consistent with research which shows the failure of ECRs in STEMM disciplines to take up training opportunities (6,27). It seems the value of such training is often under appreciated by ECRs and taking worktime for training is not encouraged by their superiors. In response to the poor uptake of modules, the CREATE training focus has moved away from on-line learning and towards provision of more presentation opportunities, networking, mentoring and other professional development. The modules remain accessible to CREATE ECRs to complete on demand.

## Discussion

### Achievements

The CREATE program continues to evolve to meet the needs of emerging pulmonary fibrosis researchers and to encourage engagement. The team is aiming to establish it as a sustainable model which will continue beyond the life of the CRE-PF funding. Feedback has been regularly sought from CREATE ECRs and others involved in the program (for example, mentors). Concerns have been addressed as they have been raised, with a flexible, evidence-based approach to building capacity in the field of pulmonary fibrosis research.

The CREATE program has guided the development of the supported researchers, thus making a major contribution to the future of pulmonary fibrosis research in Australia. During the life of the program, CREATE ECRs have been active in a range of activities important for their research careers. Some of these achievements reported by CREATE ECRs are summarised in Table 3 below.

**Table 3:**
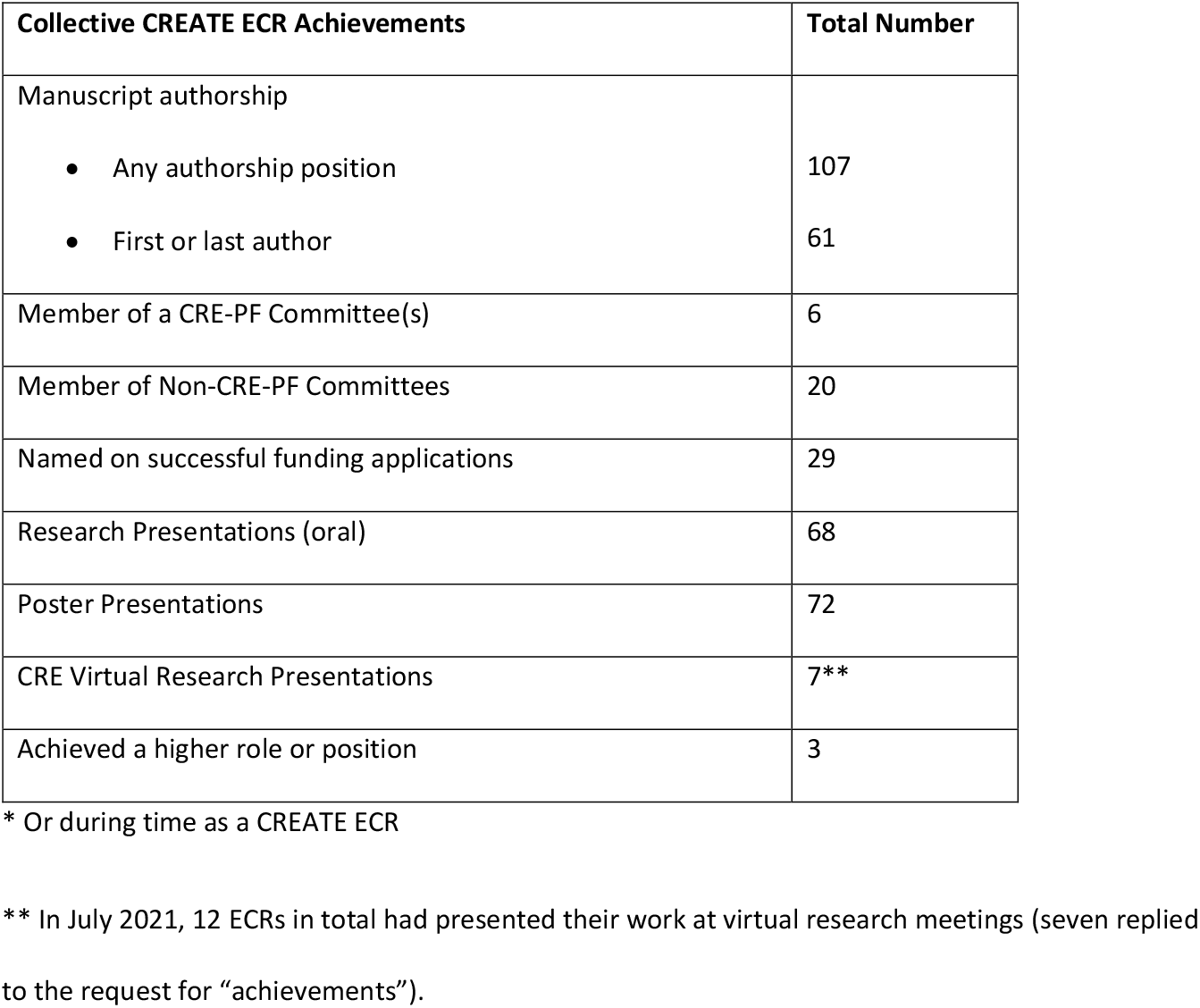
Pulmonary Fibrosis Research-related Achievements of 17 CREATE ECRs 2017-2021*.

Table 3 shows that of the 17 ECRs who responded to the request for information about their research outcomes, there have been considerable important research-related achievements by which successful research is generally measured. CREATE ECR achievements include 61 manuscripts where ECRs are named as first author or last author, 68 oral research presentations and 72 poster presentations. Contributions to the research community, as measured by membership of CRE-PF and other work-related committees, is also pleasing as it indicates both involvement and broadening of networks.

As described above in Table 3, collaboration between CREATE ECRs and between ECRs and the CRE-PF network of researchers has been actively encouraged. It is too early for published outcomes from these collaborations between ECRs, however collaborative work is underway and in each case the researchers are working towards publishable outcomes and submission of larger grant applications for their projects. CREATE ECR collaborations with other CRE-PF researchers have also been fruitful and have resulted in multiple publications. To date the work of CRE-PF investigators has produced 210 publications, at least 92 of which include one or more CREATE ECRs as an author, and there are many more yet to come. De-identified data from this brief survey can be seen in Supplementary Table 2; the full CRE publication list is available at https://www.cre-pf.org.au/cre-pf-publications.

### Challenges

Through the development of this program, difficulties have been posed by the diversity of the group, funding limitations, very busy schedules and the COVID-19 pandemic. The CREATE ECR group, made up of clinicians (respiratory physicians, physiotherapists and a clinical psychologist), basic scientists and epidemiologist/health economists are drawn from across Australia and from a wide range of sub-specialties. Most did not know one another or many of the CRE-PF investigators, and it has been hard to develop an identity as a cohesive group with a common purpose. In response to this challenge, we have flexibly expanded and persisted with CREATE group activities to bring ECRs together in various forums, including conference symposia and events, virtual research meetings and a WhatsApp group. Limited funding under the original CRE-PF grant necessarily constrained the activities CREATE was able to support. Leveraging funding through partnerships with other organisations and creativity in designing activities has enabled CREATE to extend and expand offerings. We have also pursued low cost but high impact activities such as modest collaboration grants which support multiple ECRs at once, conference prizes, virtual research meetings over zoom (to which we have free institutional access) and using WhatsApp to bring the group together. The very busy schedules of the both the ECRs and the CRE-PF investigators, plus the consequences of the COVID-19 pandemic, have resulted in fewer opportunities to bring everyone together. Our approach has been also to respond flexibly to this by finding different ways to communicate and network, seeking and responding to group feedback.

### Goal and plan for sustainability

The goal is that in the future CREATE will have sufficient funding to continue to offer grants and scholarships/stipends, along with professional development activities including bringing CREATE ECRs together for regular networking, facilitation of Industry engagement, for development of novel funding streams and collaboration building opportunities. It is clear there are many benefits to be gained by facilitating collaborative research within the wider CRE-PF network.

Funding is being targeted from a range of government and industry sources, often with the collaborative assistance of peak bodies LFA and TSANZ. In order to maintain the CREATE program into the future the CRE-PF has successfully sought industry sponsorship. Further funding for CREATE will be included in future grant applications for the CRE-PF.

## Conclusion

The CREATE Program has achieved its aim of establishing a national training scheme for respiratory researchers. In spite of a very limited budget, wide geographic distribution of participants and the multi-disciplinary nature of the cohort, we have succeeded in providing a unique, supportive academic development environment for CREATE ECRs. The willing involvement of all participants, whether senior investigators or brand-new PhD students, has provided great encouragement to these ECRs in a safe and supportive environment.

It has been important for the CREATE program to adopt a flexible and responsive approach to developing these emerging researchers at a critical time in their careers. The program provides funding and professional development opportunities within a supportive and collaborative group environment. At the completion of their training, within their institutions and from the CREATE program, all the CREATE ECRs will have an enhanced skillset which sets them up for success and longevity as a pulmonary fibrosis researcher. The CREATE program equips ECRs to effectively communicate with the scientific community and general public; work effectively and efficiently within team environments; write successful grant applications; establish and effectively lead a research group; and understand the ethical and funding framework for health research in Australia. Should they stay within the CREATE umbrella, they will gain additional skills to effectively mentor the next generation of translational scientists. The extensive research achievements to date of the CREATE ECRs as a group provide ample evidence of the benefits of the funding, mentoring and development events which have been made available to them within the program.

In the process of developing this program, there have been many lessons learned which are transferrable to other academic disciplines. In order for this type of program to succeed, we have found that it is critical to leverage and strategically distribute funding, seek and respond to feedback from the participants you are aiming to support, be flexible and nimble in response to changing international circumstances and group needs and to invest resources into building networks to support ECRs into the future.

## List of Abbreviations

CREATE: (<u>CRE A</u>dvanced <u>T</u>raining <u>E</u>nvironment)
CRE-PF: Centre of Research Excellence in Pulmonary Fibrosis
ECR: early-career researcher
LFA: Lung Foundation Australia
PAC: Program Advisory Committee
PF: pulmonary fibrosis
STEMM: science, technology, engineering, mathematics and medicine
TSANZ: Thoracic Society of Australia and New Zealand

## Declarations

### Ethics Approval and consent to participate

This manuscript reports the establishment of an education program and, for the purposes of the University of Sydney Human Research Ethics Committee (the HREC), comes under the definition of quality assurance and evaluation, not “human research” as defined in National Statement on Ethical Conduct of Human Research (29). The HREC considers reporting this research does not require ethical review; a letter has been provided to this effect. In addition,

### Consent for Publication

The authors have received written permission to use their (anonymised) feedback regarding the program in this manuscript.

### Availability of data and materials

Not applicable.

### Authors’ contributions

All authors are members of the CREATE Program Advisory Committee and have been actively involved in the design and management of the program described in this manuscript, as well as preparation of the manuscript itself.

KC and AH-C: manuscript conceptualisation, program design and management, writing original draft, critical revisions of manuscript, approval of final manuscript.

TC, PR, NG, JJ, AT, LT: program design and management, critical revisions of manuscript, final approval of the version to be published.

All authors have read and approved the final manuscript.

## Acknowledgements

The authors acknowledge, with thanks, the role of all CREATE ECR Fellows, mentors, CRE-PF investigators and staff involved in this program, including previous members of the PAC (in alphabetical order by surname): Jennifer Alison, Hayley Barnes, Christina Begka, Kaj Blokland, Chris Brereton, Mark Brooke, Janette Burgess, Amy Cashmore, Dan Chambers, Kate Christian, Britt Clynick, Tamera Corte, Ingrid Cox, Leona Dowman, Jun Fukihara, Archana Gaikwad, Ian Glaspole, Laura Glenn, Nicole Goh, Phil Hansbro, Alison Hey-Cunningham, Mariana Hoffman Barbosa, Anne Holland, Giulia Iacono, Helen Jo, Edmund Lau, Jade Jaffar, Adelle Jee, Darryl Knight, Yet Khor, Joanna Lee, Gang Liu, John Mackintosh, Jennifer Mann, Ben Marsland, Tylah Miles, Yuben Moodley, Steve Mutsaers, Vidya Navaratnam, Matthew Parker, Andrew Palmer, Ashleigh Philp, Cecilia Prele, Jyotika Prasad, Jane Read, Paul Reynolds, Debra Sandford, Claudia Sim, Alastair Stewart, Alan Teoh, Claire Thomson, Lauren Troy, Jennifer Walsh, Haydn Walters, Glen Westall, Margaret Wilsher, Hari Wimaleswaran and Qiang Zheng.

## Author Notes

Dr Katherine Christian is the CREATE Training Program Coordinator for the NHMRC Centre of Research Excellence in Pulmonary Fibrosis, Australia and a member of the Program Advisory Committee of the CREATE Program.

Dr Alison Hey-Cunningham is the Project Manager of the NHMRC Centre of Research Excellence in Pulmonary Fibrosis, Australia; staff member of The University of Sydney Central Clinical School, Sydney, Australia; and a member of the Program Advisory Committee of the CREATE Program

A/Prof Tamera Corte is a Consultant Respiratory Physician at Royal Prince Alfred Hospital, Sydney; Associate Professor at The University of Sydney Central Clinical School, Sydney, Australia; Chief Investigator A of the NHMRC Centre of Research Excellence in Pulmonary Fibrosis, Australia and a member of the Program Advisory Committee of the CREATE Program.

A/Prof Nicole Goh is a Respiratory and Sleep Consultant at Austin Health and Alfred Health, Melbourne, Australia; Clinical Associate Professor, Faculty of Medicine, The University of Melbourne; an Associate Investigator of the NHMRC Centre of Research Excellence in Pulmonary Fibrosis, Australia; and a member of the Program Advisory Committee of the CREATE Program.

Dr Jade Jaffar is Research Fellow in the Department of Allergy, Immunology and Respiratory Medicine, The Alfred Hospital, Melbourne, Australia; a CREATE Fellow of the NHMRC Centre of Research Excellence in Pulmonary Fibrosis, Australia; and a member of the Program Advisory Committee of the CREATE Program.

Professor Paul Reynolds is a Consultant Physician in the Department of Thoracic Medicine, Director of the Lung Research Laboratory at Royal Adelaide Hospital, Adelaide, Australia; a Chief Investigator of the NHMRC Centre of Research Excellence in Pulmonary Fibrosis, Australia; and Chair of the Program Advisory Committee of the CREATE Program.

Dr Alan Teoh is a Consultant Respiratory Physician at Royal Prince Alfred Hospital, Sydney, Australia; a CREATE Fellow of the NHMRC Centre of Research Excellence in Pulmonary Fibrosis, Australia; and a member of the Program Advisory Committee of the CREATE Program.

Dr Lauren Troy is a Consultant Respiratory Physician at Royal Prince Alfred Hospital, Sydney, Australia and a member of the Program Advisory Committee of the CREATE Program, NHMRC Centre of Research Excellence in Pulmonary Fibrosis, Australia.

---

**Supplementary Table 1.**
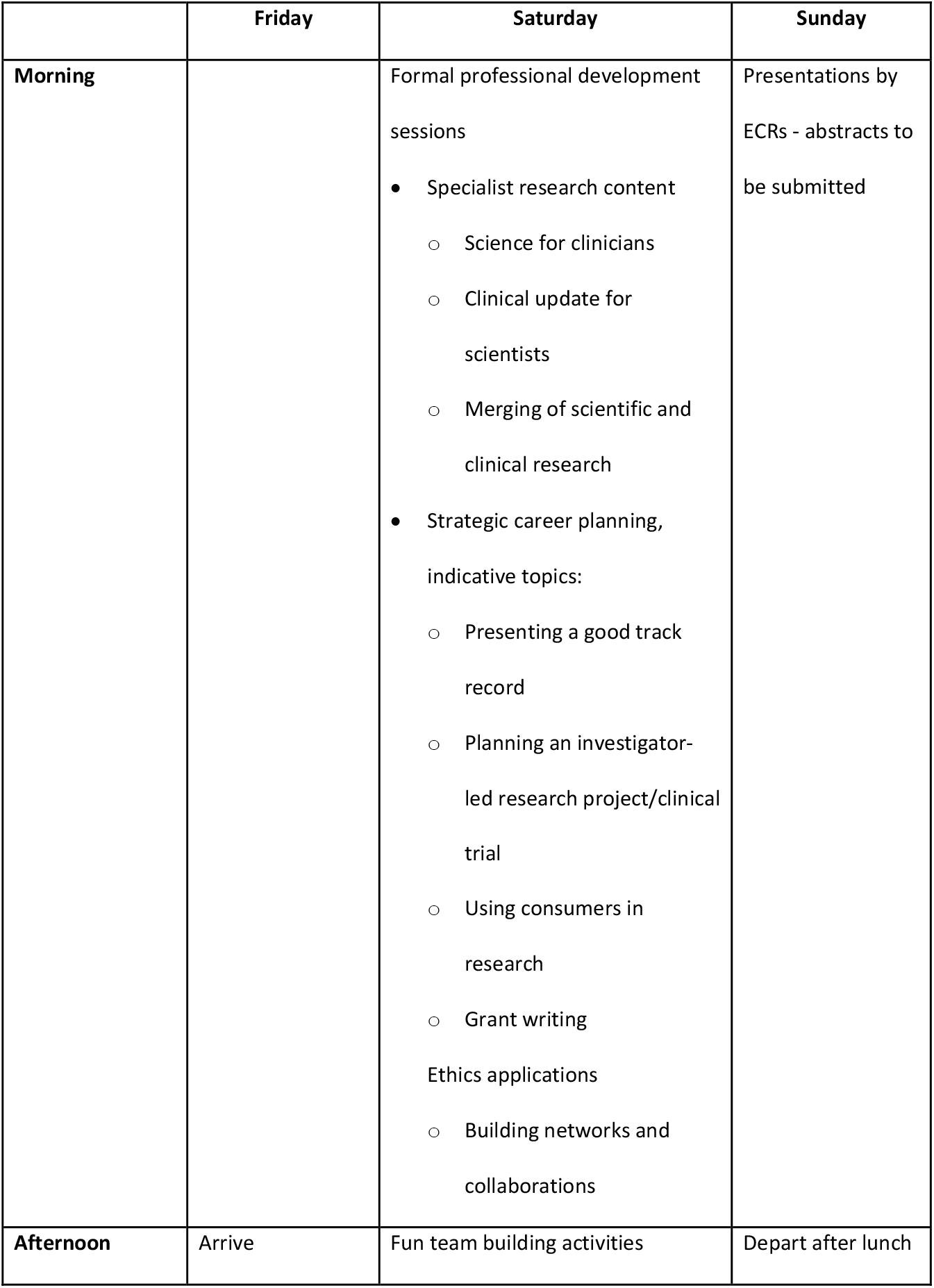
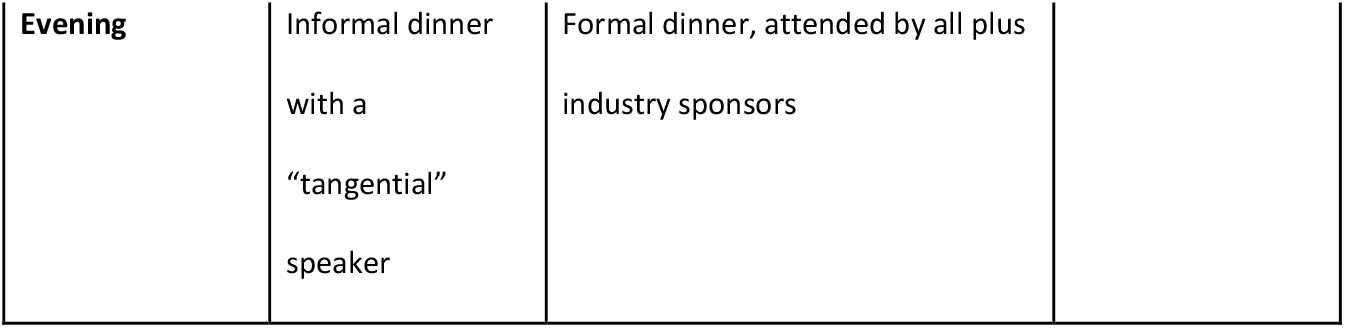
CREATE Professional Development Weekend – Program Overview.

**Supplementary Table 2.** Responses to Survey by CREATE Fellows.

|  | # manuscripts all positions | # Manuscripts 1<sup>st</sup> or last author | #CRE Committees | # Non-CRE Committees | successful funding applications | # Presentations | # Posters | CRE Virtual Presentation | Higher position | Other |
| --- | --- | --- | --- | --- | --- | --- | --- | --- | --- | --- |
| Fellow 1 | 20 | 13. F1 reported 17 | 1 | 2 | 5 | 9 | 22 | 1 |  |  |
| Fellow 2 |  |  |  |  |  |  |  |  |  |  |
| Fellow 3 | 7 | 2 + 2 in progress. Count 2 | 0 | 1 | 1 | 4 | 3 | 0 |  |  |
| Fellow 4 |  |  |  |  |  |  |  |  |  |  |
| Fellow 5 |  |  |  |  |  |  |  |  |  |  |
| Fellow 6 | 3 | 3. F6 reported 2 | 0 | 2 | 1 | 4 | 3 | 1 | NA | NA |
| Fellow 7 | 9 F7 reported 7 | 4. F7 reported 2 | 0 | 0 | 4 | 5 | 2 | 0 | 0 | NA |
| Fellow 8 |  |  |  |  |  |  |  |  |  |  |
| Fellow 9 |  |  |  |  |  |  |  |  |  |  |
| Fellow 10 |  |  |  |  |  |  |  |  |  |  |
| Fellow 11 | 7 | 6. F11 reported 7 | 0 | 0 | 1 | 2 | 1 | 0 | 0 | NA |
| Fellow 12 |  | 1 in progress | 0 | 0 | 0 | 3 | 2 |  |  |  |
|  |  | Count 0 |  |  |  |  |  |  |  |  |
| Fellow 13 | 12 | 2. F13 reported 3 | 1 | 2 | 2 | 8 | 8 | 1 | senior scientist, Grade 3 | attracted AUD\$395,020 in funding |
| Fellow 14 |  | 6. F14 reported 4 |  | 1 | 3 | 8 | 5 | 1 |  |  |
| Fellow 15 | 21 | 16. F15 reported 5 | 2 | 1 | 4 | 7 | 7 | 1 | Senior Lecturer |  |
| Fellow 16 | 5 | 1 + 1 under review | 1 TEDS | 0 | 2 scholarships | 1 upcoming | 2 | 0 | 0 | started PhD program |
|  |  | Count 1 |  |  |  |  |  |  |  |  |
| Fellow 17 | 8 | 3 | 1 | 5 | 1 | 1 | 0 | 0 | 0 |  |
| Fellow 18 | 3 | 8 BUT ONLY 1 CRE |  |  | 1 | 2 |  | 1 |  |  |
| Fellow 19 | 8 | 2 | 0 | 2 | 0 | 4 | 6 | NA | NA | Completed Fellowship training, Overseas Clinical Fellowship in ILD |
| Fellow 20 | 2 | 1 |  | 2 | 1 | 2 | 5 | 1 |  |  |
| Fellow 21 | 2 | 1 F21 reported 12, but 2 CRE |  | 2 | PhD scholarship and a previous fellow scholarship | 5 | 3 |  | Clinical lecturer |  |
| Fellow 22 |  |  |  |  |  |  |  |  |  |  |
| Fellow 23 |  |  |  |  |  |  |  |  |  |  |
| Fellow 24 |  |  |  |  |  |  |  |  |  |  |
| Fellow 25 | 0 | 0 F25 reported 1 (conf abst) | 0 | 0 | 1 | 4 | 2 | 0 | 0 | NA |
| Fellow 26 |  |  |  |  |  |  |  |  |  |  |
| Fellow 27 |  |  |  |  |  |  |  |  |  |  |
| Fellow 28 |  |  |  |  |  |  |  |  |  |  |
| Fellow 29 |  | 0 | 0 | 0 | 0 | 0 | 1 | 0 | 0 | NA |
| Total from 17 | 107 | 61 after changes F1 F6 F7 F11 F14 F15 F21 F25 F13 | 5 | 15 | 28 | 67 | 72 | 7 | 3 |  |

Note: Fellows were asked to report their number of publications "relating to the CRE". This request was interpreted differently, (and understandably) by individuals - some too high, some too low. Numbers actually counted were checked against and ultimately taken from the full list of CRE publications. The full list of CRE publications can be seen at https://www.cre-pf.org.au/cre-pf-publications

## References

1. Eley DS, Jensen C, Thomas R, Benham H. What will it take? Pathways, time and funding: Australian medical students’ perspective on clinician-scientist training. BMC Med Educ. 2017 Dec 8;17(1):242.

2. Noble K, Owens J, André F, Bakhoum SF, Loi S, Reinhardt HC, et al. Securing the future of the clinician-scientist. Nature Cancer. 2020 Feb;1(2):139–41.

3. Eley DS. The clinician-scientist track: an approach addressing Australia’s need for a pathway to train its future clinical academic workforce. BMC Medical Education. 2018 Oct 3;18(1):227.

4. Butler D. Translational research: Crossing the valley of death. Nature. 2008 Jun;453(7197):840–2.

5. Sutherland KA. Holistic academic development: Is it time to think more broadly about the academic development project? International Journal for Academic Development. 2018 Oct 2;23(4):261–73.

6. Christian K, Johnstone C, Larkins J, Wright W, Doran MR. Research culture: A survey of early-career researchers in Australia. Rodgers P, Deathridge J, Lijek RS, Rolf H, Hussain T, editors. eLife. 2021 Jan 11;10:e60613.

7. Wellcome Trust. What researchers think about the culture they work in [Internet]. Wellcome Trust; 2020 [cited 2020 Jan 17]. Available from: https://wellcome.org/reports/what-researchers-think-about-research-culture

8. Shine J, Bradlow H, O’Reilly S, Godfrey B. Women in STEM decadal plan [Internet]. Australian Academy of Science; 2019 p. 62. Available from: https://www.science.org.au/support/analysis/decadal-plans-science/women-in-stem-decadal-plan

9. Browning L, Thompson K, Dawson D. It takes a village to raise an ECR: Organisational strategies for building successful academic research careers. International Journal for Researcher Development [Internet]. 2016;7(2). Available from: www.emeraldinsight.com/2048-8696.htm

10. Browning L, Thompson K, Dawson D. Developing future research leaders: Designing early career researcher programs to enhance track record. International Journal for Researcher Development. 2014;5(2):123–34.

11. Gascoigne T. Career support for researchers: understanding needs and developing a best practice approach. Australian Council of Learned Academies Commissioned by the Department of Industry, Innovation, Science, Research and Tertiary Education; 2012 Nov.

12. Sutherland KA. Chapter 8: Resources, Training and Support for Early Career Academics. In: Early Career Academics in New Zealand: Challenges and Prospects in Comparative Perspective. Springer; 2018.

13. Department of Industry S. STEM Equity Monitor: Research workforce and grant funding data [Internet]. STEM Equity Monitor. Department of Industry, Science, Energy and Resources; 2020 [cited 2020 Apr 6]. Available from: https://www.industry.gov.au/data-and-publications/stem-equity-monitor/workforce/research-workforce-and-grant-funding-data

14. Pannell JL, Dencer-Brown AM, Greening SS, Hume EA, Jarvis RM, Mathieu C, et al. An early career perspective on encouraging collaborative and interdisciplinary research in ecology. Ecosphere. 2019;10(10):e02899.

15. Climie RE, Wu JHY, Calkin AC, Chapman N, Inglis SC, Colafella KMM, et al. Lack of Strategic Funding and Long-Term Job Security Threaten to Have Profound Effects on Cardiovascular Researcher Retention in Australia. Heart, Lung and Circulation [Internet]. 2020 Aug 14 [cited 2020 Aug 19];0(0). Available from: https://www.heartlungcirc.org/article/S1443-9506(20)30396-6/abstract

16. McKeon S, Alexander E, Frazer I, Ferris B, Brodaty H, Little M. Strategic Review of Health and Medical Research – Better Health Through Research. Canberra, Australia: Australian Government, Department of Health and Aging; 2013 Feb.

17. MacIntosh-Murray A. Poster Presentations as a Genre in Knowledge Communication: A Case Study of Forms, Norms, and Values. Science Communication. 2007 Mar 1;28(3):347–76.

18. Sutherland KA. Cultivating Connectedness and Generosity in Universities: A View of Early Career Academic Experiences in Aotearoa, New Zealand. On Education Journal for Research and Debate [Internet]. 2018 [cited 2019 Sep 23]; Available from:https://www.oneducation.net/no-03_december-2018/cultivating-connectedness-and-generosity-in-universities-a-view-of-early-career-academic-experience-in-aotearoa-new-zealand/

19. Denecke D, Feaster K. Professional Development: Shaping Effective Programs for STEM Graduate Students. Washington, DC: Council of Graduate Schools; 2017.

20. LERU. Delivering talent: Careers of researchers inside and outside academia LERU Position Paper 2018 [Internet]. Belgium: LERU; 2018 Jun. Available from: www.leru.org

21. Lee A. Nature’s Guide to Mentors. Nature. 2007 Jun 14;447:791–7.

22. Woolston C. Postdoctoral mentorship key to career success. Nature. 2019 Jan 24;565:667–667.

23. Christian K, Scott CL. Establishing the Cure Cancer Australia Mentoring Program 2017: Setting up a Successful Mentoring Program Might Be Easier than You Think. Journal of Intergenerational Relationships. 2019 Aug 16;0(0):1–10.

24. Vassallo A, Frost S, Georgousakis M. Franklin Women Mentoring Program 2017 Evaluation [Internet]. Franklin Women; 2020 [cited 2020 Nov 8]. Available from: https://franklinwomen.com.au/wp-content/uploads/2020/02/Mentoring-Program-2017-Evaluation-SummaryVFINAL.pdf

25. Bhakta D, Boeren E. Training needs of early career researchers in research-intensive universities. Intl Jnl for Researcher Dev. 2016 May 9;7(1):84–102.

26. Bogle D, Compton-Daw E, Elvidge L, Gavaghan D, Henstock J, Jones D, et al. Review of the Concordat to support the career development of researchers. 2018.

27. Christian K. Challenges faced by early-career researchers in the sciences in Australia and the consequent effect of those challenges on their careers : a mixed methods project [Internet] [Doctor of Philosophy]. [Ballarat, Victoria, Australia]: Federation University Australia; 2021 [cited 2021 Jun 17]. Available from: https://researchonline.federation.edu.au/vital/access/manager/Repository/vital:15093

28. University of Sydney. University of Sydney Research Code of Conduct 2019. 2019.

29. National Health and Medical Research Council. National Statement on Ethical Conduct in Human Research (2007) - Updated 2018 [Internet]. Commonwealth of Australia, Canberra; 2018 [cited 2019 Oct 2]. Available from: https://www.nhmrc.gov.au/about-us/publications/national-statement-ethical-conduct-human-research-2007-updated-2018

